# Gene network simulations provide testable predictions for the molecular domestication syndrome

**DOI:** 10.1101/2021.03.19.436202

**Authors:** Ewen Burban, Maud I Tenaillon, Arnaud Le Rouzic

## Abstract

The domestication of plant species lead to repeatable morphological evolution, often referred to as the *phenotypic domestication syndrome*. Domestication is also associated with important genomic changes, such as the loss of genetic diversity compared to adequately large wild populations, and modifications of gene expression patterns. Here, we explored theoretically the effect of a domestication-like scenario on the evolution of gene regulatory networks. We ran population genetics simulations in which individuals were featured by their genotype (an interaction matrix encoding a gene regulatory network) and their gene expressions, representing the phenotypic level. Our domestication scenario included a population bottleneck and a selection switch mimicking human-mediated directional and canalizing selection, i.e., change in the optimal gene expression level and selection towards more stable expression across environments. We showed that domestication profoundly alters genetic architectures. Based on four examples of plant domestication scenarios, our simulations predict (i) a drop in neutral allelic diversity, (ii) a change in gene expression variance that depends upon the domestication scenario, (iii) transient maladaptive plasticity, (iv) a deep rewiring of the gene regulatory networks, with a trend towards gain of regulatory interactions, and (v) a global increase in the genetic correlations among gene expressions, with a loss of modularity in the resulting coexpression patterns and in the underlying networks. We provide empirically testable predictions on the differences of genetic architectures between wild and domesticated forms. The characterization of such systematic evolutionary changes in the genetic architecture of traits contributes to define a *molecular domestication syndrome*.

## INTRODUCTION

Domestication is a process of rapid evolution over successive generations of anthropogenic selection, leading to adaptation to habitats created by humans and acquisition of profitable traits for them. Such innovations originate from genetic and plastic variation sustaining phenotypic shifts in domesticates compared to their wild counterparts (Fuller *et al*. 2010). In plants, traits targeted by those shifts alter architecture (more compact morphology), life-history (loss or partial loss of seed dispersal and seed dormancy, increased synchronicity of germination and ripening), as well as production- and usage-related traits (taste, increase of harvestable organs). They are often associated with convergent phenotypic changes across species (Larson *et al*. 2014), and collectively referred to as the *phenotypic domestication syndrome*.

The discovery of the genetic bases underlying variation of domesticated traits has been the focus of ample empirical work. Dozens of domestication genes have been discovered, most of which are transcription factors (Martínez-Ainsworth and Tenaillon 2016; Fernie and Yan 2019) embedded into complex gene regulatory networks (GRNs). Perhaps the most emblematic example is provided by the *Tb1* gene, which together with other genes controls maize branching architecture via hormone and sugar signaling (Doebley *et al*. 1997; Whipple *et al*. 2011; Dong *et al*. 2017, 2019). It is responsible for the strong apical dominance phenotype, i.e. repression of axillary bud outgrowth (Clark *et al*. 2006). Interestingly, in contrast to the maize allele, the *Tb1* allele from its wild ancestor (teosinte) confers a responsiveness to light when introgressed into a maize background (Lukens and Doebley 1999). It therefore appears that domestication has triggered the selection of a constitutive shade avoidance phenotype in maize (Studer *et al*. 2017), that has translated into a loss of phenotypic plasticity. Along this line, recent results indicate that reduced Genotype-by-Environment (GxE) interactions may be a general consequence for traits targeted by human selection. For example, genomic regions displaying footprints of selection explain less variability for yield GxE than “neutral” regions (Gage *et al*. 2017). Decreased phenotypic plasticity during domestication likely results both from the stability of human-made compared with wild habitats and from selection for stable crop performance across environments; it has yet to be characterized in other crops and for a broader range of traits.

In addition to the *phenotypic domestication syndrome*, genome-wide sequencing data have revealed the outlines of a *molecular domestication syndrome*. This molecular syndrome includes a loss of genetic diversity through linked selection and constriction of population size due to sampling effects (Yamasaki *et al*. 2005). Severity of those genetic bottlenecks as estimated by nucleotide diversity loss, ranges from 17% to 49% in annuals while often no loss is observed in perennial fruit crops (reviewed in Gaut *et al*. 2015). The combined effect of bottlenecks, increased inbreeding (Glémin and Bataillon 2009) and linked selection in domesticates translates into shrink in effective population size, which in turn reduces the efficacy of selection against deleterious mutations (Moyers *et al*. 2018). Although increased recombination rate in domesticates compared with their wild relatives may partially compensate this effect (Ross-Ibarra 2004), fixation of deleterious mutations in domesticates and a resulting genetic load is often observed as exemplified in African rice (Nabholz *et al*. 2014), grapevine (Zhou *et al*. 2017), and maize (Wang *et al*. 2017).

Regarding molecular phenotypes assessed by transcriptomic surveys, data are still scarce and emerging patterns not as clear. Measures of variation of gene expression in domesticates relative to their wild counterparts either reveal a significant loss, as in rice, cotton (Liu *et al*. 2019), beans (Bellucci *et al*. 2014); a significant gain as in tomato (Sauvage *et al*. 2017); or no substantial change as in soybean (Liu *et al*. 2019), olives (Gros-Balthazard *et al*. 2019), and maize (Swanson-Wagner *et al*. 2012). In the latter, however, reduced variation in expression was observed at domestication candidate genes, indicating that selection primarily acts on *cis*-acting regulatory variants (Hufford *et al*. 2012): most evolutionary-relevant mutations affecting the evolution of gene expressions are located in (or in the close vicinity) of the domestication genes. This result was further confirmed in F1 hybrids from maize / teosinte crosses where large differences in expression were primarily caused by *cis*-divergence, and correlated with genes targeted by selection during domestication (Lemmon *et al*. 2014).

Beyond quantitative measures of gene expression, domestication is also associated with gene network rewiring. A pioneer work in maize indeed indicates that 6% of all genes display altered co-expression profiles among which, genes targeted by selection during domestication and/or breeding are over-represented (Swanson-Wagner *et al*. 2012). Interestingly, networks encompassing domestication targets display greater connectivity in wild than in domesticated forms as if selection had triggered connection loss to/from these genes. In beans, coexpression networks at the genome level revealed a global excess of strong correlations in domesticates compared with wild, the latter being sparser with more isolated nodes and smallest connected components than the former (Bellucci *et al*. 2014). In contrast to maize, little qualitative difference was reported as for networks surrounding selected and neutral contigs.

While population genetic tools have been broadly used to estimate domestication bottlenecks and associated genetic load in plants (Eyre-Walker *et al*. 1998; Tenaillon *et al*. 2004; Wright *et al*. 2005; Gaut *et al*. 2015; Kono *et al*. 2016; Liu *et al*. 2017; Wang *et al*. 2017), a theoretical framework that considers *molecular domestication syndrome* as a whole allowing to make predictions beyond verbal models is still in its infancy (Stetter *et al*. 2018). Here, we propose to simulate the evolution of gene regulatory networks in a population submitted to domestication-like pressures. We used a modified version of a classical gene network model (the ‘Wagner’ model, after Wagner 1994, 1996) to represent the complex genetic architecture of gene expression regulation, and tracked the evolution of genetic diversity, of gene expression plasticity, and of network topology in scenarios featuring (i) a temporary drop in the population size (bottleneck), and (ii) a substantial change in the selection regime. The default demographic scenario was defined based on maize, an outcrosser crop with a relatively simple domestication (a single origin for the crop with a moderate domestication bottleneck); we further studied alternative domestication scenarios (African rice, peal millet, and tomato) to assess the robustness of our conclusions. Simulations aim at providing a general framework to explore a multitude of scenarios and life-history traits, and experimentally testable predictions.

## MATERIALS AND METHODS

### Gene network model

The gene network model was directly inspired from Wagner (1996), with minor changes detailed below. An illustration of a simplified (3 genes) network evolution under this model is given Figure 1. Individual genotypes were stored as *n* × *n* interaction matrices ***W***, representing the strength and the direction of regulatory interactions between *n* transcription factors or regulatory genes. All genes have the potential to regulate other genes of the network (although such feedback is not mandatory). Each element of the matrix ***W_ij_*** stands for the effect of gene *j* on the expression of gene *i*; interactions can be positive (transcription activation), negative (inhibition), or zero (no direct regulation). Each line of the ***W*** matrix can be interpreted as an allele, i.e., the set of regulatory sites in the promoter of the gene. The model considered discrete regulatory time steps, and the expression of the *n* genes, stored in a vector ***P***, changes during the development of an individual as ***P*_*t*+1_**=*F*(***W P_t_***), where *F* (*x*_1_, *x*_2_, …, *x_n_*) applies a sigmoid scaling function *f*(*x*) to all elements to ensure that gene expression ranges between 0 (no expression) and 1 (full expression). We used an asymmetric scaling function as in Rünneburger and Le Rouzic (2016); Odorico *et al*. (2018): *f* (*x*) = 1 /(1 + λ*e^−μx^*), with *λ* = (1 – *a*)/*a* and *μ* = 1/*a*(1 – *a*). This function is defined such that *a*=0.2 stands for the constitutive expression (in absence of regulation, all genes are expressed to 20% of their maximal expression).

**Figure 1:**
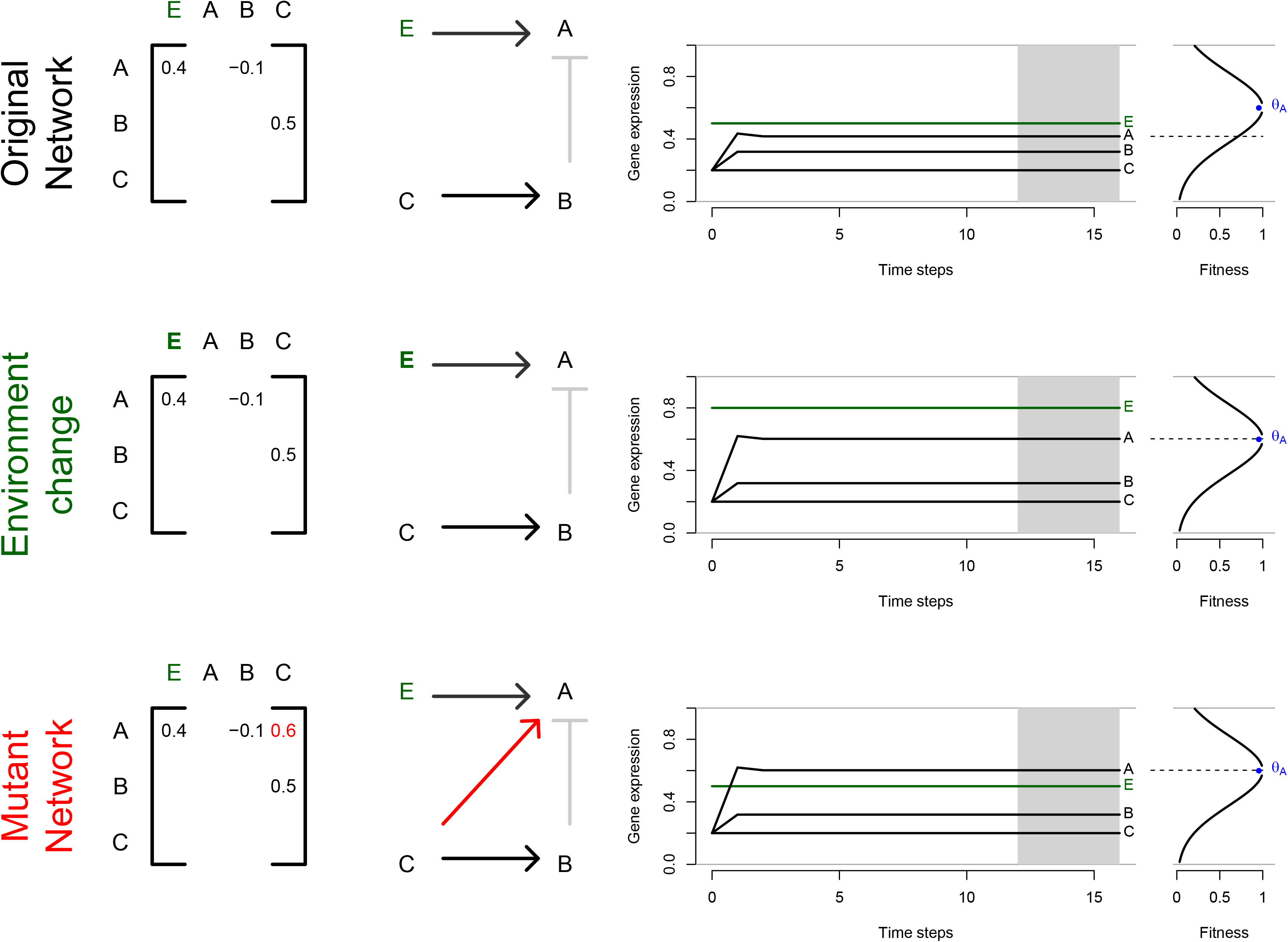
Gene network model. An original network with three genes (A, B, C) and an environmental sensor gene (E) is illustrated. The genotype of an individual is provided by the matrix of interactions (left panel), which provides the strength and direction (middle panel) of regulatory interactions: A is a plastic gene influenced by E (and B), B is up-regulated by C but its expression does not depend on the environment, C is not regulated and therefore expressed constitutively (level of expression set to 0.2). This example assumes that selection targets only the expression of gene A, which optimum is *θ_A_*=0.6 independently from the environment. Top row; expression of E is set to 0.5. The kinetics of the network during 16 time-steps shows a rapid stabilization after 3 steps, the expression of the genotype is however distant from the expression target which represents a fitness cost (fitness<1). Middle row: Upon an environmental change (expression of E changes from 0.5 to 0.8), the expression of the genotype reaches the expression target, and the fitness is maximal. Bottom row: Without environmental change, maximum fitness can be reached with a mutant network whereby C now regulates A; such a mutant network would have a fitness advantage over the original network in this environment. Variance of expression are computed for each gene on the four last time-steps (shaded area), and networks which have not reached a stable state at that point (non-null variance) suffer a fitness penalty.

The kinetics of the gene network was simulated for 16 time-steps in each individual, starting from ***P*_0_**=(*a*, …,*a*). The simulation program reports, for each gene *i*, the mean 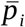 and the variance *V_i_* of its expression level over the four last time steps. A non-null variance characterizes networks that have not reached equilibrium at 16-4=12 time-steps, either because of slow network dynamics or because the network is unstable (cyclic pattern). In addition to this traditional framework, we considered that one of the network genes was a “sensor” gene influenced by the environment. This makes it possible for the network to react to an environmental signal, and evolve expression plasticity. In practice, the environmental signal at generation *g* was drawn in a uniform distribution *e_g_* ~ *U*(0,1) and the value of the sensor gene was *e_g_* at each time step (the sensor gene had no regulator and was not influenced by the internal state of the network).

### Population model

The gene network model was coupled with a traditional individual-based population genetics model. Individuals were diploids and hermaphrodites, and generations were non-overlapping. Reproduction consisted in drawing, for each of the *N* offspring, one (in case of selfing) or two (outcrossing) parents randomly with a probability proportional to their fitness. Parents gave two gametes, a gamete containing a random allele at each of the *n* loci (assuming free recombination). There was no recombination between regulatory sites at a given locus (the model assumes *cis*-regulation only, so that all regulatory sites are close to the gene). The genotype of an individual was defined by both inherited gametes; the ***W*** matrix from which the expression phenotype was calculated was obtained by averaging out maternal and paternal haplotypes. In the ‘Wagner’ model, regulatory effects are additive; the regulatory effects of both alleles average out, and the effects of transcription factors add up. Yet, even if regulatory effects are additive, the mapping between the strength of regulation and gene expression is non-linear (sigmoid). As a consequence, the model accounts for both dominance and epistasis at the gene expression level, strong regulators (activators or inhibitors) being dominant over weak regulators. For instance, the gene expression in a loss-of-function heterozygote will be closer to the functional homozygote than to the mutant homozygote.

Individual fitness *w* was calculated as the product of two components, *w* = *w_U_*×*w_S_*. The first term *w_U_* corresponds to the penalty for networks that have not reached stability, 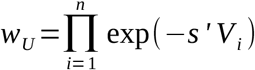, *s*′ being the strength of selection on gene expression variance (i.e., selection against expression instability). The second term *w_S_* corresponds to a Gaussian stabilizing selection component, which depends on the distance between the expression phenotype and a selection target 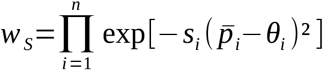, *s_i_* standing for the strength of stabilizing selection on gene *i*. As detailed below, some genes were not selected (in which case *s_i_* = 0), some genes were selected for a stable optimum *θ_i_* (“stable” genes), while a last set of genes were selected for optima that changed at every generation *g* (“plastic genes”), half of them being selected for *θ_ig_*=*e_g_*, and the other half for *θ_ig_*=1 – *e_g_*. Selection was moderate (*s*=10) for most simulations, albeit stronger selection (*s*=50) was also tested (Figure S1).

Mutations occurred during gametogenesis with a rate *m*, expressed as the mutation probability per haploid genome. A mutation consists in replacing a random element of the ***W*** matrix by a new value drawn in a Gaussian distribution centered on the former value 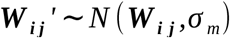, where *σ_m_* is the standard deviation of mutational effects. In this model, mutations affect gene regulatory regions only (i.e., protein sequences do not evolve); mutations occurring in the promoter of a gene affects primarily its own expression, but the rest of the network may also be affected when this gene regulates other ‘downstream’ genes (Figure 1).

### Domestication scenario and parameterization

Domestication was associated with two independent changes in the simulation parameters: a temporary demographic bottleneck (decrease in population size), and a change in the gene expression optima (directional selection). In order to calibrate simulations with realistic parameters, we used simplified versions of documented domestication scenarios. The default scenario features a protracted model of maize-like domestication involving a moderate bottleneck starting about 9,000 years (generations) ago with a bottleneck strength of *k*=2 *N_b_*/*T_b_* = 2.45 (Wright *et al*. 2005), *N_b_* and *T_b_* being the effective population size and the duration of the bottleneck. Simulations were thus split in three stages: (i) a long “burn-in” stage (*T_a_*=12,000 generations in the largest population size *N_a_*=20,000 that was computationally tractable, unless specified otherwise) aiming to simulate predomestication conditions, after which the “ancestral” species is expected to harbor genotypes adapted to wild conditions (selection optima *θ_a_*, drawn in a uniform *U* (0,1) distribution at the beginning of each simulation for “stable” genes, fluctuating optima for “plastic” genes), (ii) a bottleneck of *T_b_*=2,800 generations (Eyre-Walker *et al*. 1998), during which the population size was reduced to *N_b_* =3,430 individuals, and selection optima switched to *θ_b_*, and (iii) *T_c_* = 6,200 generations of expansion of the domesticated species (population size to *N_c_*=20,000), while the selection optima remained to the “domestication” conditions *θ_b_*. Selection under dometication conditions, compared to ancestral condition, implied more stable genes and less plastic genes. For computational feasibility, the regulation network size was limited to 24 genes (+1 environmental signal), from which 12 were under direct selection. Before domestication, the network encompassed 12 unselected, 6 stable, and 6 plastic genes (Figure S2). At the onset of domestication, we modified the selection regime to mimic increased environmental stability and, in turn, decreased plasticity (12 unselected, 10 stable and 2 plastic genes, Figure S2). The mutation rate was set to *m* = 10^−3^/ gamete/ generation, which, given the estimated mutational target of 24 genes of 1kb (average estimated length of enhancers from Oka *et al*. 2017; Ricci *et al*. 2019) roughly corresponded to a per-base mutation rate of 3×10^−8^ par generation, close to the maize estimate (Clark *et al*. 2004).

In addition to the maize default domestication scenario, we considered three additional domestication scenarios (African rice, pearl millet, and tomato). Only the demography (timing and strength of the bottleneck) was modified; the strength and mode of selection before and after domestication was identical to the maize scenario. Unknown demographic parameters were replaced by educated guesses as detailed below, and the maximum population size was capped at N=20,000. The domestication of the African rice (*Oryza glaberrima*) is characterized by a long bottleneck (k=0.61), starting 10,000 generations ago and lasting 8200 generations (Cubry et al 2018). Although the domestication might have started later than the beginning of the bottleneck, we considered that the selection switch occurred simultaneously with the bottleneck. The African rice is a selfer, the selfing rate was set to 0.98. The domestication of the pearl millet (*Cenchrus americanus* syn *Pennisetum glaucum*) is more recent (4800 generations ago), with a short bottleneck (k=1.89) (Clotault et al 2021, Burgarella et al 2018). Just like maize, pearl millet is an outcrosser. Finally, the tomato (*Solanum lycopersicum*) was domesticated about 6400 years ago, with a long (5800 generations) bottleneck (k=0.17) (Arnoux et al 2020). The ancestral species was featured by a low diversity (estimated *N_a_*=1,600), in contrast to the other large-ancestral population cases. The tomato is a selfer (selfing rate 0.98). Scenario parameters are summarized in Table 1. In addition to the four default domestication scenarios (maize, African rice, pearl millet, tomato) described above, we explored control simulations to disentangle the contribution of the bottleneck and the selection switch in emerging patterns, based on the maize scenario (‘Default’): a scenario with no bottleneck, and a scenario with no selection switch. Given the importance of these three scenarios (Default, no bottleneck, no selection switch) in the analysis of the results, simulations were run with a longer burn-in (*T_a_*=24,000) to ensure that the network was close to the mutation-selection-drift equilibrium at the onset of domestication. We further assessed the sensitivity of our results for the maize default domestication scenario to independent changes in parameters values by (1) increasing the number of genes of the GRN, from 24 to 48, and doubling the number of selected genes and the mutation rate per genome accordingly; (2) setting the mutation rate to 0 at the time of domestication to evaluate selection response from standing variation only; (3) modulating selection intensity both through a decrease in selected genes count (by two-fold), and through a modification of the fitness function to simulate stronger selection; (4) dissociating selection switch from a loss of plasticity, either by maintaining the selection for plasticity over genes during domestication, or by keeping the same number of plastic genes before and after domestication; (5) testing the effect of a harsher (10 times less individuals) bottleneck. We also assessed the sensitivity of the model to arbitrary parameters influencing the gene network dynamics, such as the number of time steps, or the selection on network instability. All scenarios were replicated 1000 times; unless specified otherwise, the reported variables were averaged over all individuals from the population; figures report the grand mean over the replicates; colored areas stand for the 10% - 90% quantiles over the simulation replicates.

**Table 1:** Demographic parameters associated with all four domestication scenarios.

|  | Maize | African rice | Pearl millet | Tomato |
| --- | --- | --- | --- | --- |
| Burn-in $T_a$ | 24,000 | 12,000 | 12,000 | 12,000 |
| Burn-in $N_a$ | 20,000 | 20,000 | 20,000 | 1,600 |
| Bottleneck $T_b$ | 2,800 | 8,200 | 900 | 5,800 |
| Bottleneck $N_b$ | 3,430 | 2,500 | 850 | 490 |
| Bottleneck k | 2.45 | 0.61 | 1.89 | 0.17 |
| After $T_c$ | 6,200 | 1,800 | 3,900 | 600 |
| After $N_c$ | 20,000 | 20,000 | 5,400 | 20,000 |
| Selfing rate | 0 | 0.98 | 0 | 0.98 |

### Model output and descriptive statistics

For each simulation run, summary statistics were computed every 100 generations. The output includes the population mean and variance of (i) the absolute fitness *w*, (ii) gene expressions 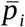, (iii) gene regulations ***W_ij_*** for all pairs of genes. In addition, the environmental index *e_g_* and all selection optima *θ_g_* were recorded. Effective population sizes were estimated as 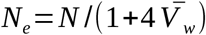 (Walsh & Lynch 2018), where 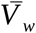 stands for the variance in the relative fitness 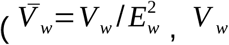, and *E_w_* being the population variance and the population mean of the absolute fitness, respectively). When computed over a time interval (e.g. over the duration of the bottleneck), the harmonic mean effective size 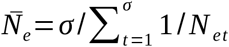 was reported.

A proxy for neutral molecular variance around gene *i* was obtained by reporting the average population variance of the ***W_ij_*** for a subset of genes *j* which expression was very low (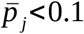 over the whole simulation), as regulatory sites sensitive to nonexpressed transcription factors are expected to evolve neutrally.

Environmental reaction norms (gene expression plasticity) were estimated for each gene *i* by regressing the average expression 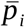 over the environmental index *e_g_*, taken over a sliding window of 10 consecutive measurements (1000 generations).

The effect of gene regulations ***W_ij_*** being quantitative (and thus, never exactly 0), the presence/absence of a connection in the network was determined by the following procedure: the expression phenotypes ***P*** and 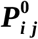 were calculated both from the full ***W*** matrix, and from each of the *n*^2^ possible 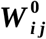 matrices in which ***W_ij_*** was replaced by 0. The regulation ***W_ij_*** was considered as a meaningful connection when the Euclidean distance 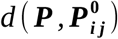 exceeded an arbitrary threshold of 0.1. Using other thresholds shifted the number of connections upward or downward, but did not affect the results qualitatively.

Genetic correlation matrices were estimated directly from the population covariances in gene expressions (hereafter called ***G*** matrices, although they reflect here all genetic components and not only additive (co)variances). The evolution of ***G*** matrices was tracked by computing the distance between consecutive matrices ***G_g_*** and ***G*_*g*+500_** in the simulation output. In practice, genetic covariances were turned into genetic correlation matrices, and then into genetic distance matrices 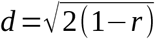. The difference between both genetic distance matrices was calculated from their element-wise correlation, as in a Mantel test (function mantel.rtest in the R package ade4, Dray and Dufour 2007). Network topological features, including the number of clusters used as an index of modularity, were measured with the package igraph (Csardi and Nepusz 2006).

### Implementation and Data availability

The simulation model was implemented in C++ and compiled with gcc v-7.5.0. The simulation software is available at https://github.com/lerouzic/simevolv. Simulation runs were automated via bash scripts, and simulation results were analyzed with R version 4.0 (R Core Team 2020). All scripts (simulation launcher, data analysis, and figure generation) are available at https://github.com/lerouzic/domestication.

Supplementary figures are available from the publisher web site.

## RESULTS

We used a gene network model encompassing 24 transcription factors to simulate the *molecular domestication syndrome* and provide testable predictions regarding (i) adaptation and the evolution of plasticity, (ii) the evolution of molecular and expression variance and (iii) the extent of network rewiring. Regulation strength between genes was modelled as a quantitative variable directly affected by mutation at regulatory sites, so that individual genotypes were stored in a matrix of interactions among all genes (Figure 1). Our simulations featured plants undergoing a rather classic protracted domestication scenario with a single bottleneck. Demographic parameters were inspired by four documented domestication histories, two outcrossers (maize and pearl millet) and two selfers (African rice and tomato). We modeled the selection switch associated with domestication both as a change in the gene expression optima and a partial loss of plastic responses. The default maize domestication scenario was compared with simulations without bottleneck (albeit a selection switch), and simulations without selection switch (albeit a bottleneck), and we also explored independent variation of parameters values to explore the sensitivity of our results. The whole simulation approach is summarized in Figure S3.

### Adaptation during habitat shift

The strong selection switch resulted in an immediate change in absolute fitness which dropped to < 0.1%, mimicking transient fitness loss of wild plants during habitat shift — a wild individual would have a probability < 0.001 to be selected by a breeder over a modern crop strain (Figure 2). Fitness was slowly regained as domesticated plants adapted to their new cultivated habitat. Fitness recovery was slower in the Default scenario including a bottleneck, the end of which was featured by an increase in the rate of fitness gain. With the maize default scenario, the population has entirely recovered its initial fitness roughly 9,000 generations after the selection switch, the process being 2000 generations faster in absence of a bottleneck (Figure 2). Most of the evolutionary change was due to new mutations, as simulations without mutations from the beginning of domestication, i.e., adapting from the standing genetic variation only, did show a very limited response to selection (Figure S4 B).

**Figure 2:**
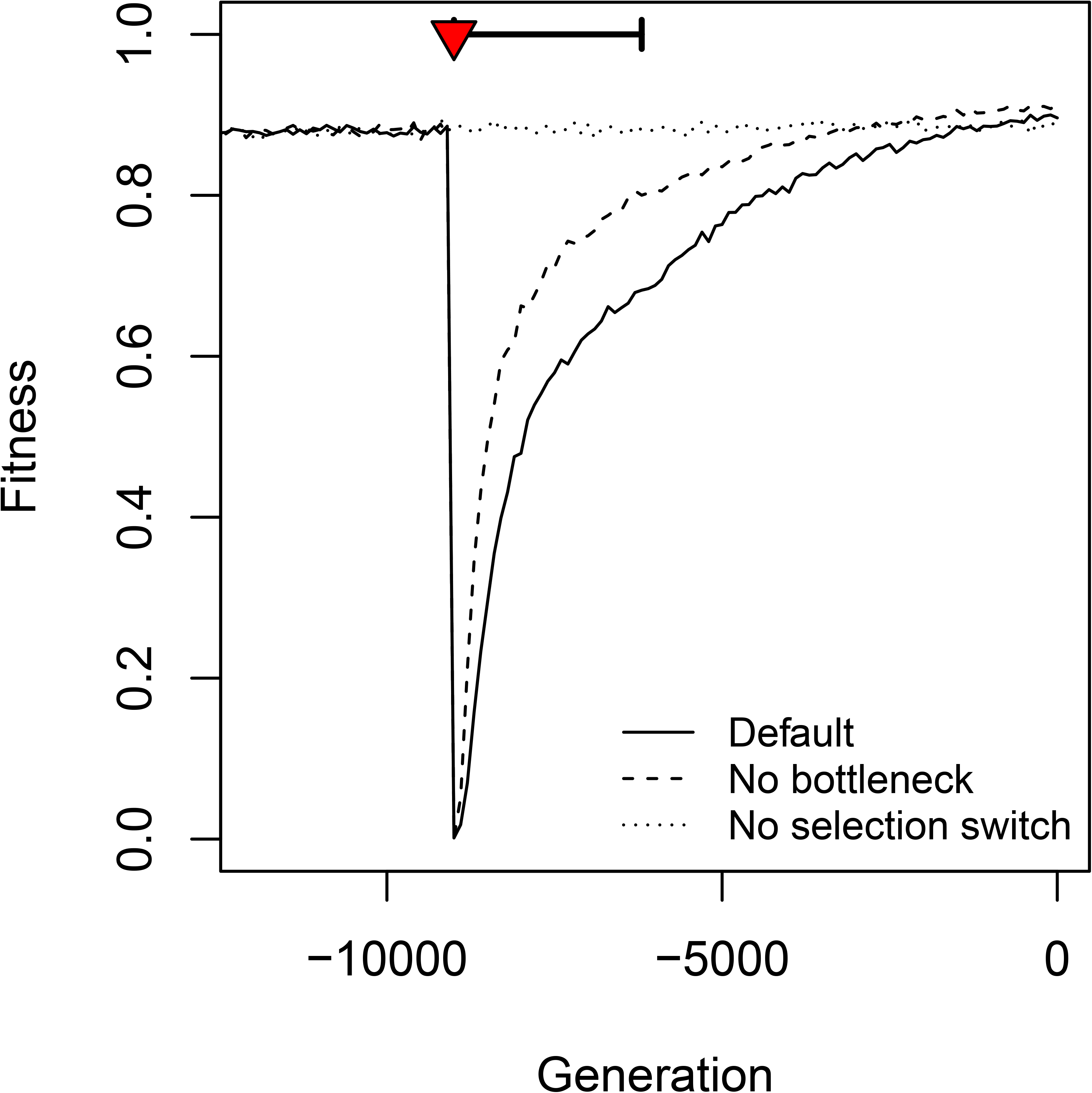
Effect of domestication on the average population fitness. The fitness drop corresponds to the switch in the optimal gene expressions at the onset of domestication (selection switch, red triangle). The population bottleneck is indicated as a thick horizontal segment. Three scenarios were considered: a full maize domestication scenario with selection switch and bottleneck (Default, plain line); a scenario without bottleneck (hyphenated line); a scenario with a bottleneck but constant selection (no switch, dotted line). The figure shows the average over 1000 simulations for each scenario.

We simulated the loss of plasticity during domestication as a change in selection regime for four plastic genes (out of 6) towards stable selection or neutrality (Figure S2). The speed at which the gene network evolved increased by a factor of roughly 13 when the selection regime shifted (Figure 3A). The selection switch translated into an abrupt change in reaction norm for genes that became selected for a flat reaction norm (plastic → stable in Figure 3B). We indeed observed a rapid loss of plasticity, showing that it was an evolvable feature that responded to selection. More surprisingly, however, the loss of plasticity also affected genes that (i) were no longer under direct selection (Plastic → Neutral in Figure 3B), and (ii) were supposed to remain plastic (Plastic → Plastic in Figure 3B), albeit to a lower extent. This shortterm maladaptive evolution highlighted the genetic constraints during the rewiring of the network caused by the selection switch. Immediately after it, plastic genes were still tightly connected to genes that were selected to evolve a flat reaction norm, and the first stage of this evolutionary change involved a maladaptive trade-off. It was slowly resolved by rewiring the connections across genes. Note that the bottleneck retarded slightly the evolutionary change (Figure 3A), as adaptive plasticity was recovered faster in constant population-size simulations (Plastic → Plastic Figure 3B). Maladaptive plasticity did not evolve in simulations where plastic genes were under the same selection regime before and after domestication (figure S5E), showing that it resulted from underlying constraints of the network, where selection-triggered changes in reaction norms at some genes affect the evolution of plastic genes.

**Figure 3:**
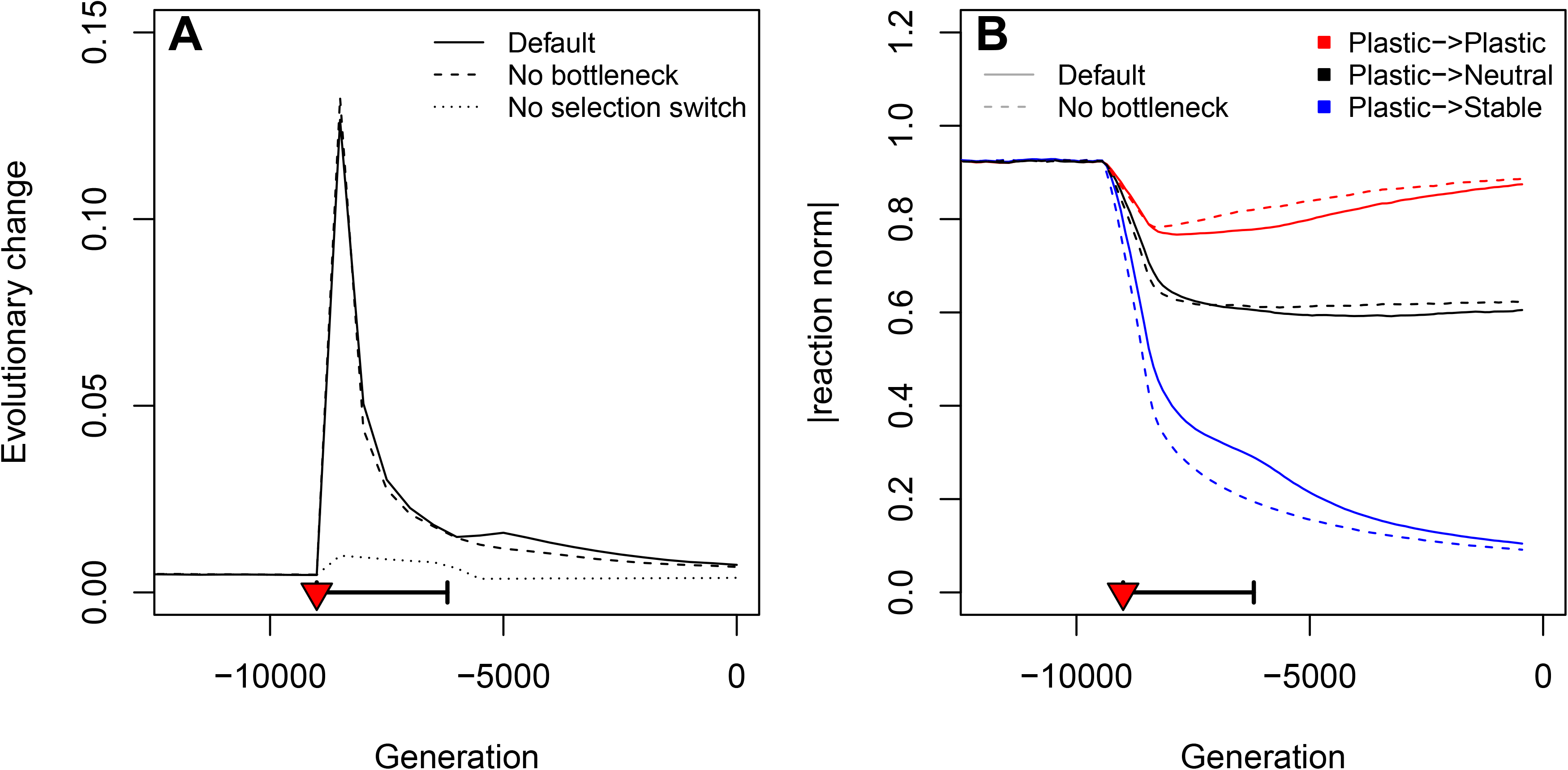
Changes in the rate of evolution and in plastic reaction norms upon domestication. A: Speed of evolutionary change in the regulatory network as measured by the Euclidean distance between the regulation strengths of successive (500 generations apart) average genotypes ***W***. Three scenarios are illustrated: full maize domestication (Default, plain lines), no bottleneck (hyphenated lines), no selection switch (dotted lines). The selection switch and population bottleneck are indicated as a red triangle and a thick horizontal segment, respectively. B: Evolution of the average reaction norm for genes that were selected to be plastic before domestication. Two scenarios are illustrated: maize full domestication (Default, plain lines), and no bottleneck (hyphenated lines). Red lines stand for genes selected to remain plastic after domestication, black lines indicate genes that were unselected in anthropic conditions, and blue lines genes that were selected to lose their expression plasticity.

### Molecular variation is affected by both demography and selection

Regulation strength was modeled as a quantitative variable directly affected by mutation (Figure 1). For any given gene in the network, we measured its neutral molecular variance among individuals of the population as the average variance of the regulation strength at regulatory sites that had no influence on gene expression. Hence, molecular variance is an analogous measure of neutral nucleotide genetic diversity of the genes of the network.

Based on empirical evidence, the first signal that we expected was a loss of neutral genetic diversity. The variance indeed dropped at the beginning of the domestication (Figure 4A). Such variance drop was driven by genetic drift, that increased during the bottleneck. The maximum observed drop in genetic diversity was ~30% loss during the bottleneck for the default scenario. Recovery was slow and still ongoing at the end of the simulations.

**Figure 4:**
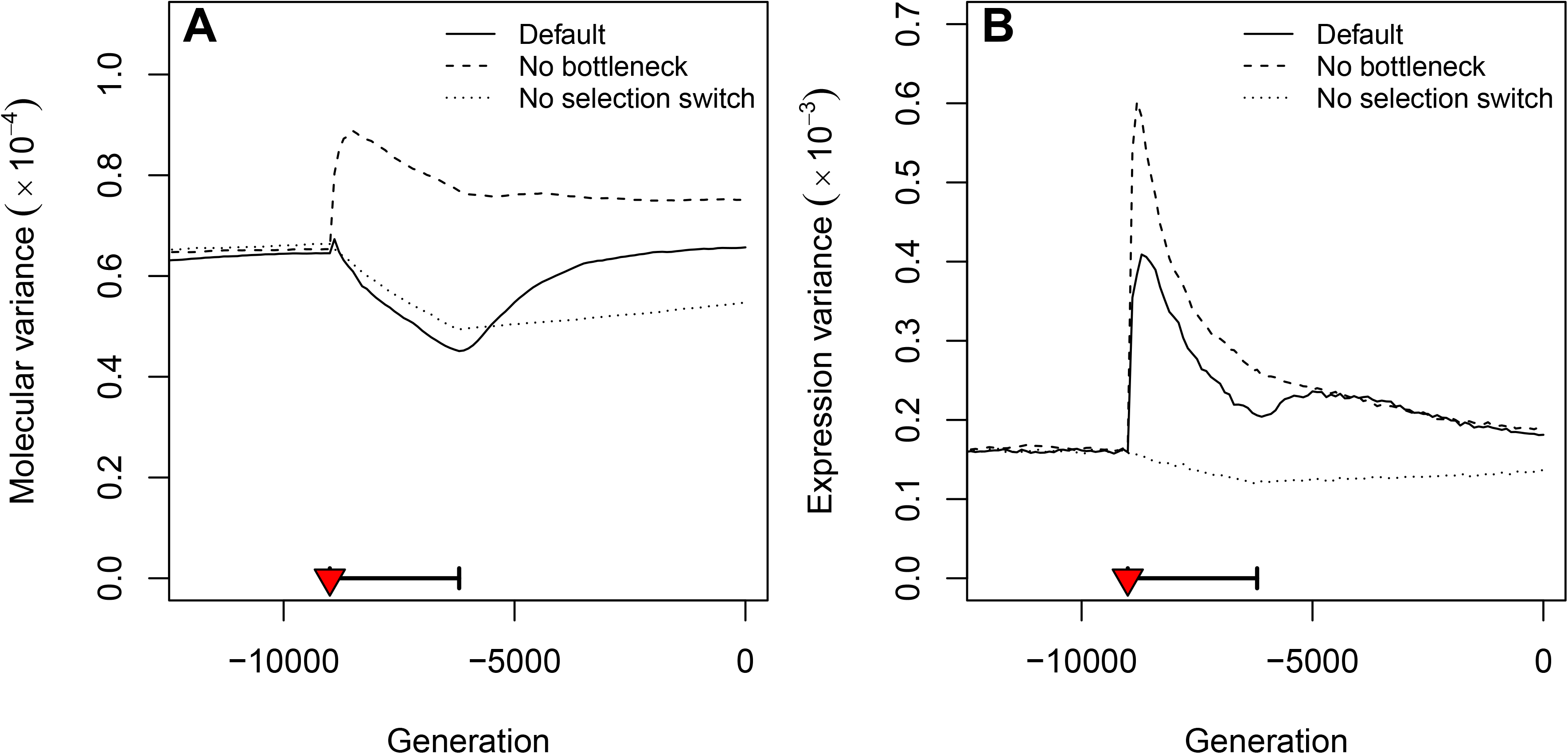
Evolution of genetic and expression variation through time. The population neutral molecular variance (A) was estimated from the regulation sensitivity to unexpressed transcription factors, which measures the genetic diversity at neutral loci that are in complete linkage disequilibrium with the network genes. The population expression variance (B) stands for the within-population “phenotypic” variance of gene expressions, averaged over all genes. The selection switch and population bottleneck are indicated as a red triangle and a thick horizontal segment, respectively. The figure shows the average over 1000 simulations for each scenario (same scenarios as in Figures 2 and 3A)

In addition to change in molecular variance, we investigated the evolution of phenotypic (expression) variance during the domestication. Our results showed that in contrast to the neutral genetic variance, phenotypic variance may increase during domestication (Figure 4B). Expression variance bursts, absent from the simulations without selection switch, can be associated with ongoing adaptation: they corresponded to the segregation of selected variants that brought the phenotype closer to the new optimum. Domestication was thus associated with an increase in the gene expression variance, as a result of the balance between the selection switch (which increased temporarily the variance, Figure 4B) and the bottleneck (which slightly reduced the variance, Figure 4B). In case of a stronger bottleneck, however, the expression diversity was reduced at the selection switch showing that the net effect on phenotypic diversity strongly depends on the details of the domestication scenario (Figure S4D).

### Domestication is associated with the rewiring of gene networks

Genetic correlation matrices (**G** matrices) were estimated from the population covariances in gene expressions. Genetic correlations evolved rapidly after domestication, and this evolution was driven both by the change in the selection regime and by the bottleneck (Figure 5A, Figure S6A). Domestication resulted in (i) a slight increase in the average coexpression from 0.11 to 0.18 (Figure 5A), and (ii) a redistribution of genetic correlations, with less distinct clusters of correlations after domestication (Figure 5B). The slight trend towards larger coexpressions results from a diversity of evolutionary changes depending on status of genes before and after domestication (Figure S6B). Overall strong correlations weakened during domestication, while many weak coexpression signals increased (Figure S6B).

**Figure 5:**
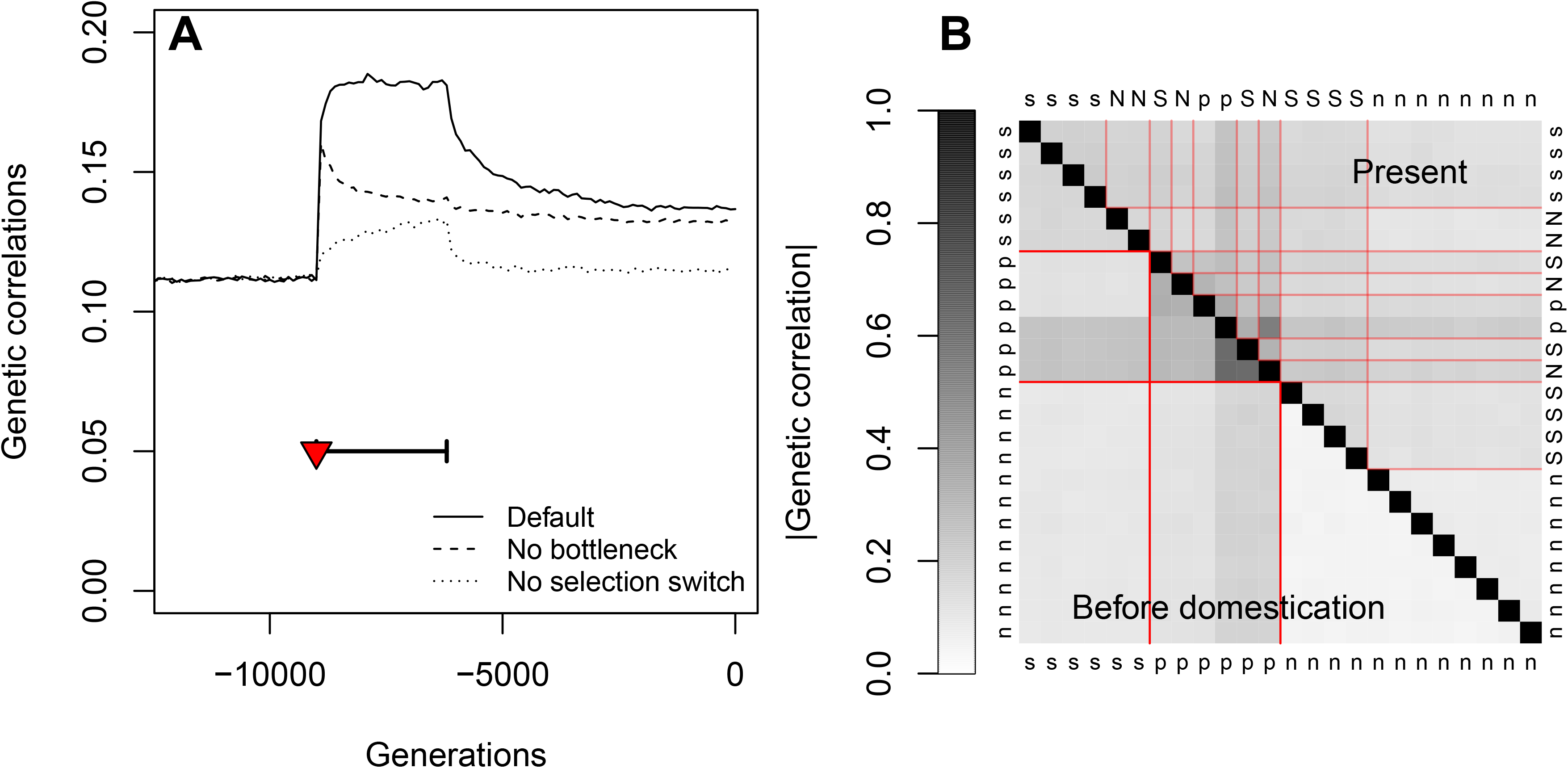
Consequences of domestication on gene coexpression. (A): Evolution of the average absolute value of within-population genetic correlations. The selection switch and population bottleneck are indicated as a red triangle and a thick horizontal segment, respectively, for the same three scenarios as in Figure 2. (B) Average genetic correlation for each pair of genes, at generation −9000 (just before the onset of domestication) below the diagonal, and at generation 0 (last generation of the simulations), above the diagonal. Gene selection status is indicated (n: nonselected, s: stable, p:plastic); capital letters indicate genes whose selection status changed during domestication. Red lines delimitates gene categories, before and after domestication. The two groups of plastic genes correspond to genes selected to correlate positively and negatively with the environmental index.

We explored the evolution of the GRN topology during domestication, by tracking the evolution of the number of connections. We observed a strong signal of network rewiring during the first stage of domestication, with an increase (by a factor > 10) of the rates of both gained and lost connections, immediately after the selection switch (Figures 6A and S7A). This rewiring was solely due to the selection switch, as there was no effect of the bottleneck alone on the network evolution. The rewiring was associated with a systematic excess of gained connections over lost connections, i.e., domestication caused an increase in the total number of connections (Figure 6A). As a consequence of the gain of new connections, the number of clusters decreased (some connections appeared between previously independent modules, Figure 6B). New connections appeared to be distributed evenly across the network (Figure S7B and S8).

**Figure 6:**
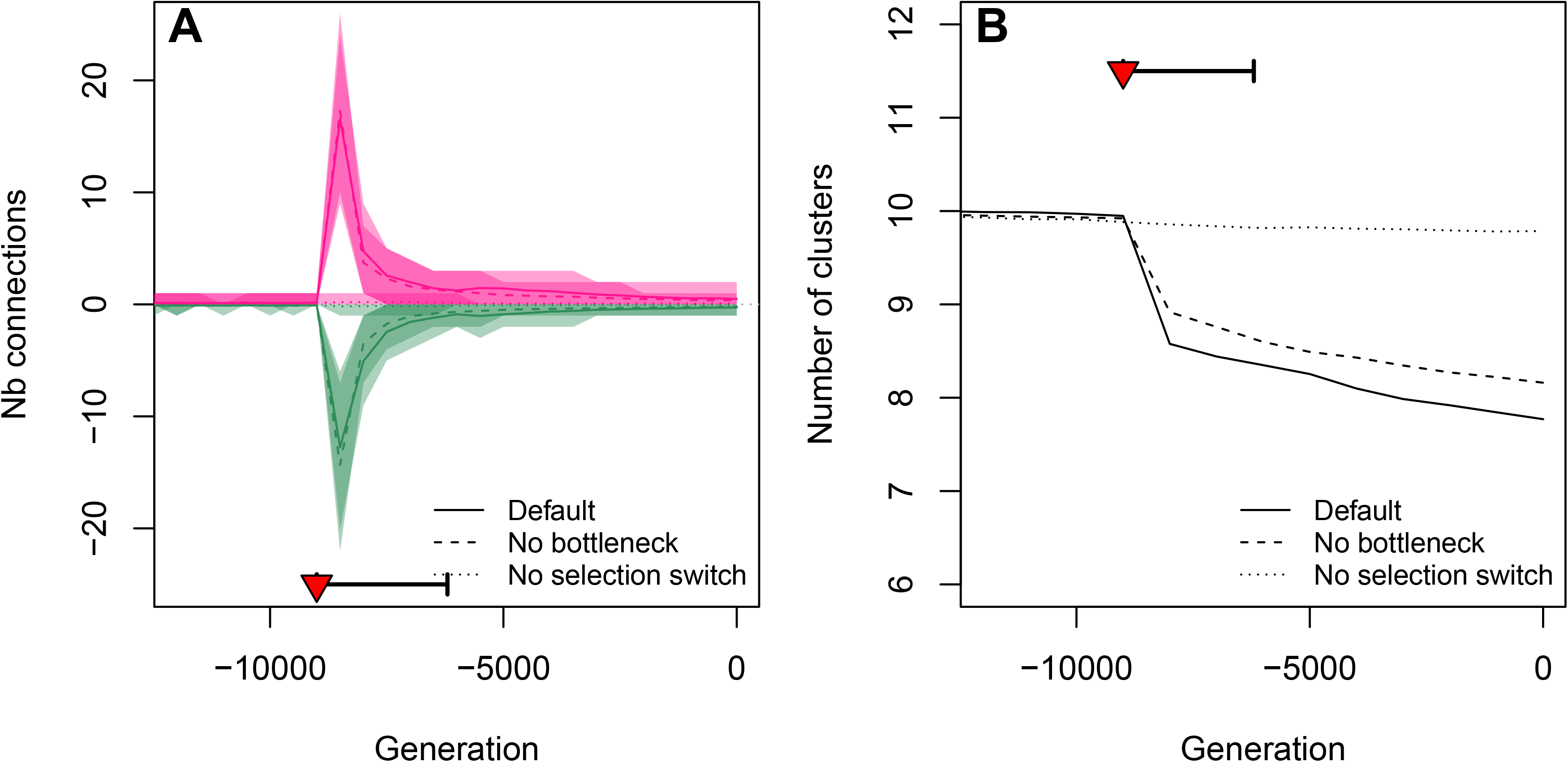
Evolution of gene network properties. The presence/absence of a regulatory connection was determined based on its effect on gene expression (see methods). A: Connection gains (pink) and losses (green) were counted over windows of 500 generations. The drop in the number of clusters (B) corresponds to new connections among existing clusters. The selection switch and population bottleneck are indicated as a red triangle and a thick horizontal segment, respectively. The three scenarios are the same as in Figure 2.

### The molecular syndrome does not depend on the domestication scenario

Based on the maize domestication scenario, we defined a list of molecular consequences of domestication, featuring (i) a drop in the molecular (genetic) variance, (ii) an increase in the phenotypic (gene expression) variance, (iii) the evolution of gene expression plasticity, with a stage during which plasticity is maladaptive, (iv) the rewiring of gene networks, with a general increase in the number of connections, and (v) a slight increase in gene expression correlations, corresponding to a loss of modularity in the underlying regulatory networks. We assessed the robustness of these results to the domestication scenario by simulating alternative demographic features, inspired by the domestication history of three plants (African rice, pearl millet, and tomato). These scenarios differ by the strength and the duration of the bottleneck (Table 1 and Figure 7), and by the rate of self-fertilization (African rice and tomato are selfers, while maize and pearl millet are outcrossers).

**Figure 7:**
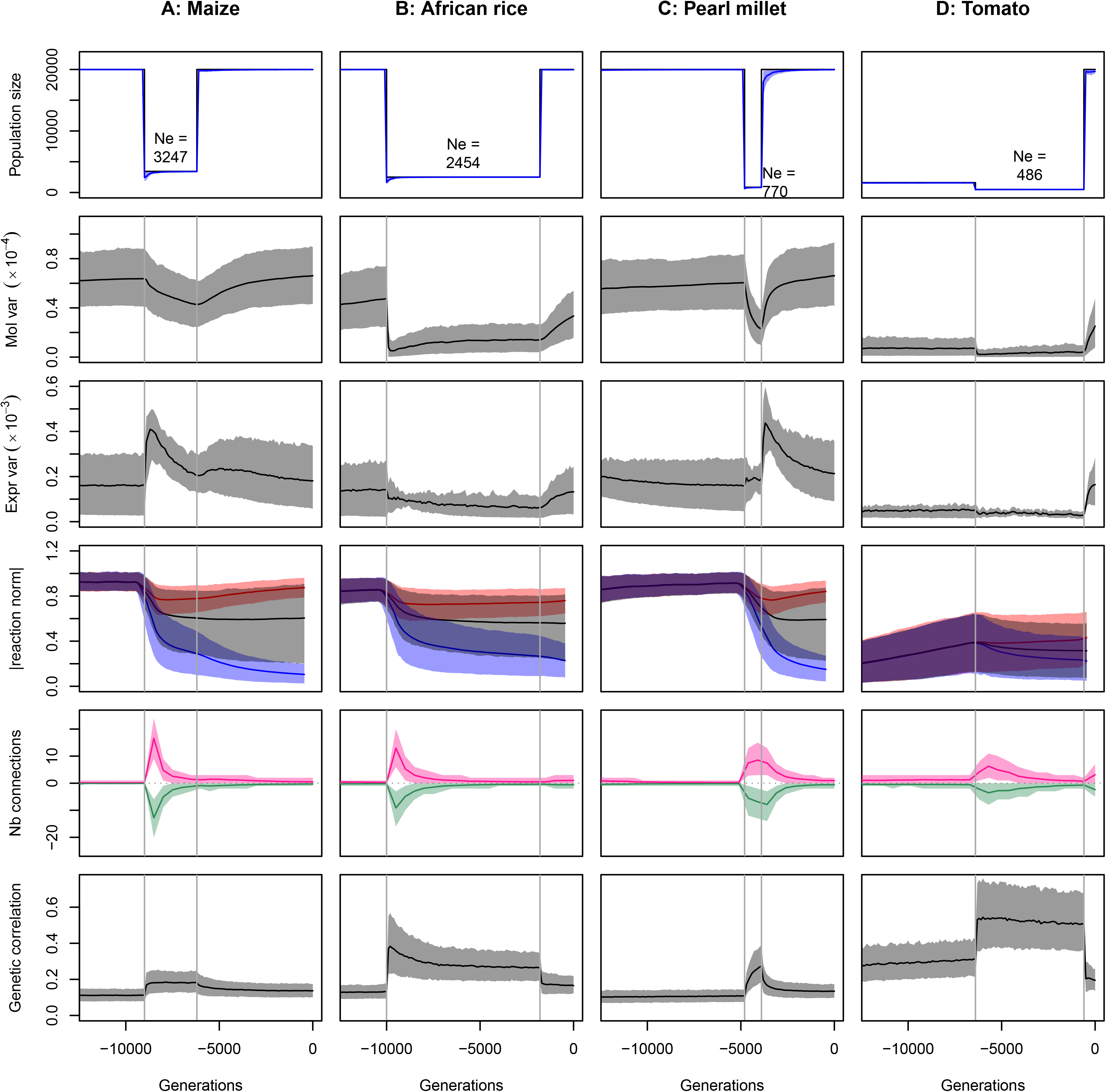
Influence of the domestication scenario on the simulation results. A: maize domestication scenario (for reference), B: African rice, C: Peal millet, D: Tomato. The four scenarios differ by the timing, strength, and duration of the bottleneck, by the demography before and after the bottleneck, and by the selfing rate (Table 1). Plain lines represent the average of each variable over 1000 simulations, shaded areas stand for the 10% - 90% quantiles. First row: census and effective population sizes. Census sizes are model parameters, effective population sizes *N_e_* were estimated from the variance in fitness (see methods). Effective population sizes indicated in the figure were computed as the harmonic mean over the whole bottleneck. Second row: Molecular variance, as in Figure 4A. Third row: Expression variance, as in Figure 4B. Fourth row: average reaction norm, as in Figure 3B. Fifth row: Number of connections, as in Figure 6A. Sixth row: Average genetic correlation, as in Figure 5A.

Overall, most of the molecular evolution observed in the maize scenario was reproducible (Figure 7). The neutral molecular variance drops during the bottleneck in all scenarios, and raises again after the bottleneck. As predicted from the maize scenario, the direction of the evolution of the variance in gene expression was sensitive to the demographic scenario, and depends on a complex balance between drift, selection, and selfing rate. The evolution of plasticity was very similar in maize, African rice, and pearl millet. Interestingly, however, the extremely small ancestral population size of tomato hampered the evolution of plasticity. In all cases, the number of network connections evolved similarly as in maize — network rewiring at the onset of domestication, with an excess of connection gains *vs*. losses. Patterns of pleiotropy in the network (average expression correlation) were consistent with an overall strong short- and mild long-term increase, except for the recent marked expansion of population size in tomato that translated into a decrease in pleiotropy below the initial level (Figure 7).

## DISCUSSION

Domestication is a complex process, involving deep modifications of the demographic, environmental, and selective context in which populations evolve. Here, we explored the consequences of domestication-like changes on the evolution of gene regulatory networks underlying domestication traits, combining a population bottleneck, directional selection and phenotypic canalization, simulated as the evolution of selection pressure towards decreased plasticity i.e., environmental stability of phenotypes.

### Adaptive dynamics under domestication

We observed that the bottleneck had a substantial effect on genetic diversity, including (i) a substantial loss of neutral genetic (molecular) diversity (Figure 4A), (ii) a moderate loss of expression variance (Figure 4B). These observations are in line with theoretical expectations. When the population size drops, genetic diversity is expected to be lost progressively, as the inbreeding coefficient increases by a factor (1–1/2*N_e_*) every generation. How much of the initial diversity of the species survives the bottleneck depends on the strength and the duration of the population size drop; in our simulations, parameterized from the maize domestication scenario, about 70% of the initial neutral diversity survived the bottleneck. This estimate matched the 60% of mean pairwise diversity retained in “neutral” maize regions as defined as those located 5 kb away from genes, with π=0.00691 and 0.0115 in maize and teosintes, respectively (Beissinger *et al*. 2016).

Less expected perhaps was the fact that even such a mild bottleneck penalized substantially the response to anthropic selection (Figure 2). This may be due to less frequent occurrence of adaptive mutations during the bottleneck or a diminished efficiency of selection, or a combination of both. In the simulations, the bottleneck was associated with a burst of segregating adaptive alleles (Figure 4B), which suggests a two-stage domestication scenario: (i) during the bottleneck, the adaptive alleles that segregate (either from the standing genetic variation and/or from new mutations) increased the population fitness, but tend to have suboptimal effects (e.g. negative side effects on well-adapted genes are illustrated by plastic genes whose reaction norm diminishes while they are continuously selected to be plastic, Figure 3B); (ii) after the end of the bottleneck, a new set of adaptive alleles can invade the population (because more mutations are available and selection is more efficient), fine-tuning genetic effects, e.g. on reaction norms (Figure 3B). Hence, we expect mutations segregating during the first stage and surviving to drift to display greater effects than those segregating during the second stage. In line with this prediction, early work on maize domestication has identified several quantitative trait loci (QTLs) with large effects, some of which were fine-mapped down to individual genes such as *Tb1* (Doebley *et al*. 1997; Studer *et al*. 2011) and *Tga1* (Wang *et al*. 2005). Examples of early mutations with large effects have also been recovered in tomato (Frary *et al*. 2000), in wheat (Simons *et al*. 2006), in rice (Konishi *et al*. 2006; Li *et al*. 2006), in barley (Komatsuda *et al*. 2007) among others. These large QTLs that most likely encode early domestication targets stand as exceptions in the overall architecture of domestication traits dominated by small-effect QTLs as recently reported in maize (Chen *et al*. 2020).

Most phenotypic changes associated with domestication are controlled by mutations in transcription factors, and therefore involve a re-orchestration of gene networks (Martínez-Ainsworth and Tenaillon 2016) as described in cotton (Rapp et al. 2010), maize (Hufford et al. 2012), bean (Bellucci et al. 2014) and tomato (Sauvage et al. 2017). Consistently, in our simulations, the gene network was deeply rewired, as the rate of gain/loss connections increased by more than one order of magnitude (Figure 6A). This effect was solely due to the shift in the selection regime. Before domestication, the population was well-adapted to an arbitrary wild type fitness landscape, involving genes which expression was constant and genes which expression was selected to track the environment. The structure of the underlying network evolved so that expressions of genes of the same type were genetically correlated, suggesting direct or indirect regulatory connections. When the fitness landscape changed, some genes that were previously correlated were forced to become independent. The results suggest that this was easier to achieve by adding connections rather than removing them, illustrating evolution by genetic tinkering instead of re-engineering. Interestingly, there was no apparent cost to this additional complexity, as the fitness after domestication reached similar levels as before domestication.

### Model approximations

Gene network models based on Wagner (1994) are built on a set of simplifying assumptions: the network dynamics is discretized and simplified (e.g. no distinction between RNA products and proteins), mutations can affect gene expression only (transcription factors do not evolve), there are no interactions between transcription factors (their effects adds up), and a given transcription factor can act both as an activator and a repressor. Little is known about the potential effect of such details on the general dynamics of the network. We confirmed that the number of developmental time steps did not affect the simulation results (except if very low, < 8) (Figure S9 B and C), nor the number of time steps during which network instability was measured (Figure S9D). Selection on the network stability did not have a perceptible effect on the results (Figure S9 E).

For the sake of realism, and to connect the model results to quantitative genetics theory, we proposed several changes to the original framework from Wagner (1996). We adopted the setting used in e.g. Siegal and Bergman (2002), in which gene expression was considered as a quantitative character, with a continuous scaling function between 0 (no expression) and 1 (maximal expression), instead of the traditional on/off binary setting (Wagner 1996; Ciliberti *et al*. 2007). We used an asymmetric sigmoid scaling (as in Rünneburger and Le Rouzic 2016) to ensure that a non-regulated gene has a low constitutive expression (here, 20% of the maximal expression). Our model allows for the possibility to evolve a plastic response. We added a perfect environmental cue as an input of the network through a sensor gene, which expression was reflecting the environmental index during the whole network dynamics. The literature provides alternative settings to introduce plasticity in the Wagner model, such as the introduction of the environmental cue as the starting state of the network, mimicking developmental plasticity (Masel 2004), or trans-generational plasticity (Odorico *et al*. 2018).

Computational constraints limited the population size to a maximum of N=20,000. Estimates of the effective population sizes of both maize and teosinte vary roughly between 10^5^ and 10^6^ depending on data/methods/models (Eyre-Walker 1998; Tenaillon et al. 2004; Beissinger et al. 2016; Wang et al. 2017), suggesting that genetic drift before and after the bottleneck was substantially larger in the simulations than expected in a realistic domestication scenario. Domestication scenarios were also greatly simplified, with a single bottleneck. Refinements of this initial setting could include multiple expansion waves of semi-domesticated forms, as well as rapid population growth and gene flow with wild relatives post-domestication (Beissinger *et al*. 2016; Kistler *et al*. 2018). For simplicity, we parameterized the bottlenecks by setting the census size (*N*) to the effective population sizes (*N_e_*) documented in the literature. Because of selection, *N_e_*<*N*, which made bottleneck slightly stronger than expected. Yet, the difference was modest (< 10%, Figure 5), and was unlikely to affect the results. Larger population size would raise the neutral diversity, but is unlikely to impact general outcomes. Due to computational constraints, we also had to limit the number of generations prior to domestication (*T_a_*) for some simulations; as a consequence, “wild” populations were not necessarily at mutation-selection-drift equilibrium. However, the effect remains limited compared to the strong effects due to domestication (e.g., Figure 5A, *T_a_*=24,000, vs. Figure 5C, *T_a_*=12,000 prior to domestication).

Likewise, network size also had to be limited to n=24 genes, as the complexity of the gene network algorithm increases with the square of the number of genes. Defining a realistic size for a gene network remains problematic, as, in fine, most genes are connected through correlated regulations. Nevertheless, we considered here only transcription factors (or TF-like regulators, such as regulatory RNAs), which have the potential to affect the expression of other genes.

Finally, how selection affects the expression level of such TFs remains quite arbitrary. For simplicity, we considered stabilizing selection directly on the gene expression level — a common setting in similar studies (e.g. Siegal & Bergman, 2002). This remains an oversimplification, as the relationship between gene expression, physiological characters, life history traits and fitness can be very complex. For instance, Draghi & Whitlock (2012) mapped *n* genes into *m* traits via a *n*×*m* transition matrix, stabilizing selection being applied at the phenotypic level, translating into indirect selection on gene expressions. Yet, if the relationship between gene expression and selected phenotypes is monotonous, applying a multivariate bell-shaped fitness function on gene expression probably remains an acceptable approximation, assuming that the details of the fitness function does not affect deeply the evolution of gene networks.

In the default scenario, most of the response to selection was due to new mutations, as the standing genetic variation alone could not explain more than about 20% of the (log) fitness recovery (Figure S4). The contribution of standing genetic variation to the response to selection is a complex function of the mutational variance, the strength of stabilizing selection before domestication, and the strength of directional selection during domestication (Stetter *et al*. 2018). The simulations thus correspond to a harsh domestication scenario in this respect, where the number of selected traits and the phenotypic changes induced by domestication were both large compared to the phenotypic diversity of the wild ancestor. We also considered that the expression level of only half of the network genes was under direct selection pressure — this would happen if half of the TFs were regulating directly key enzymes or growth factors. Simulating twice less selected genes did not affect the qualitative outcomes of the model (Figure S5).

### The molecular syndrome of domestication

Simulations confirm that the domestication process is expected to be associated with several characteristic signatures (S) at the molecular level; S1: a decrease of allelic diversity, S2: a change in gene expression variance, S3: the rewiring of the gene regulatory networks, and S4: less modularity of coexpression patterns.

The loss of genetic diversity (S1) was both due to the bottleneck (genetic drift removed rare alleles from the population) and to the selection shift (selective sweeps decreased the genetic diversity at linked loci), it is thus expected to be a general signature of domestication (Figure 4A). Empirically, a loss of genetic diversity is indeed always associated with domestication, although its amplitude may vary (reviewed in Gaut *et al*. 2015).

The direction and magnitude of the evolution of gene expression variance (signature S2) depends on the balance between selection and drift; bottlenecks tend to reduce diversity, while a shift in the selection regime tends to increase it transiently (segregation of adaptive variants). Given our simulation parameters, inspired from the maize domestication scenario featuring a mild bottleneck, expression variance increased (Figure 4B). This was not necessarily the case with all parameter combinations, as a stronger bottleneck as in African rice led to a decrease in both molecular and expression variance (Figure 7). The strength and the pattern of selection also affect the speed and the nature (soft vs. hard) of the selective sweeps, which may differ across species. As a consequence, domestication is not expected to be associated with a systematic evolution of gene expression variance: it may increase when the bottleneck is moderate, as in maize, or decrease in species where the bottleneck was drastic and/or associated with an autogamous mating system, such as rice, cotton (Liu *et al*. 2019), and beans (Bellucci *et al*. 2014).

Genetic networks were rewired (signature S3) and evolved towards less modularity (Figure 5B), as a consequence of swapping the selection pattern among genes (shift in the optimal expression for stable genes, and loss of plasticity for others). The network was less plastic after domestication, which was a consequence of a modelling choice (domestication was associated with a decrease in the number of genes expected to respond to the environmental cue). New connections occurred among previously isolated modules, but former connections were not all eliminated. As a result, the rewiring of regulatory connections lead to a moderate increase in gene coexpressions (signature S4), associated with a loss of structure in the coexpression network (uncorrelated genes became correlated, and strongly correlated genes became more independent). This illustrates a realistic evolutionary scenario towards non-adaptive complexity, where the final network structure is not the more efficient one, but rather results from the accumulation of successive beneficial mutations in an existing, constrained genetic background. Empirically, we therefore predict that connections involving genes targeted by domestication should increase rather than decrease, in line with observations in beans where coexpression networks revealed a global excess of strong correlations in domesticates compared with wild (Bellucci *et al*. 2014). Global increase in genetic correlations (Figure 5A & Figure 7) should translate into greater constraints and pleiotropy, and less independent modules. Interestingly, the general increase in genetic correlations was associated with a trend towards homogenization, i.e. strong correlations tended to weaken whereas uncorrelated genes became slightly correlated (Figure 5B). Empirical comparisons at 18 domestication-related traits between two independent populations of offspring generated by the intermating of multiple parents from a teosinte population and from a maize landrace, revealed several interesting features in line with our observations: only a subset of genetic correlations (33 out of 153) were conserved between teosinte and maize, teosinte correlations were more structured among trait groups (Yang *et al*. 2019). Investigating carefully the transcriptome evolution for several pairs of domesticated / ancestral populations will be necessary to assess the predictive power of our theoretical model.

The genetic diversity available in modern cultivated species is often considered as a limitation to further response to artificial selection. Controlling recombination has been proposed as crucial for plant breeders to engineer novel allele combinations and reintroduce diversity from wild crop relatives (reviewed in Taagen *et al*. 2020). Yet, if the domestication syndrome was also associated with changes in the pleiotropy of the genetic architecture, genetic progress might also be limited by undesirable genetic correlations among traits of interest (Yang *et al*. 2019). Understanding how genetic constraints evolved under anthropic selection and whether it is possible to avoid or revert them requires a better understanding of the complex non-linear mapping between domestication genes and phenotypes.

## ACKNOWLEDGEMENTS

We are grateful to Sylvain Glémin for insightful comments on the manuscript. We thank Clémentine Vitte for useful literature suggestions on enhancers. Simulations were performed on the core cluster of the Institut Français de Bioinformatique (https://www.france-bioinformatique.fr/ifb-core/).

## FUNDING INFORMATION

This work received financial support from two grants overseen by the French National Research Agency (ANR): one from the LabEx BASC -- Biodiversité, Agroécosystèmes, Société, Climat (ANR-11-LABX-0034) to ALR, and the DomIsol project (ANR-19-CE32-0009) to MIT. We thank the GDR 3765 “Approche Interdisciplinaire de l’Évolution Moléculaire” for travel support to EB. EGCE and GQE-Le Moulon benefit from the support the Institut Diversité, Écologie et Évolution du Vivant (IDEEV), and GQE-Le Moulon from Saclay Plant Sciences-SPS (ANR-17-EUR-0007).

## Notes

https://github.com/lerouzic/domestication

### Competing Interest Statement

The authors have declared no competing interest.

