## Supplementary Figures S1 to S9. for "Gene network simulations provide testable predictions for the molecular domestication syndrome"

##### Supplementary Figure S1.

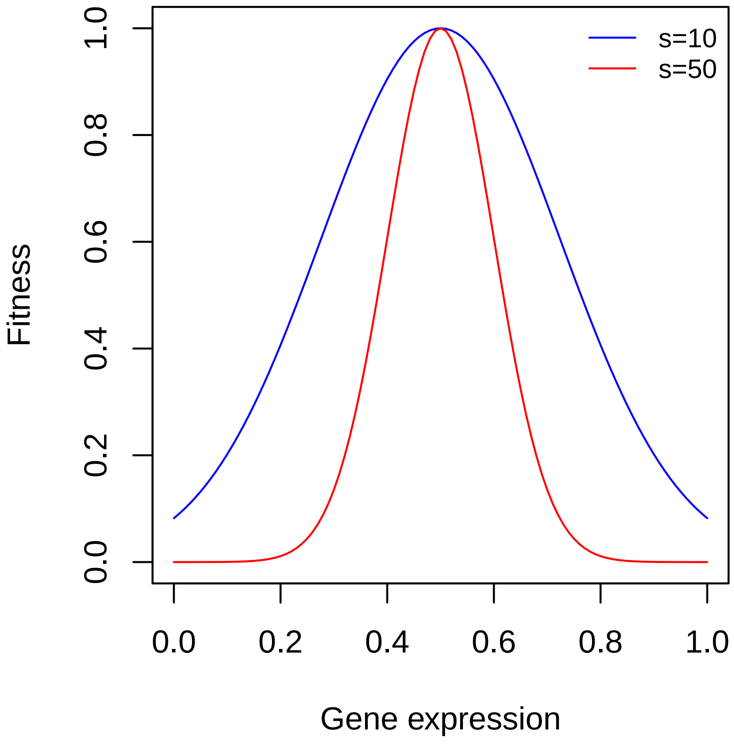


**Illustration of the fitness function.** The fitness functions is bell-shaped ($w = e^{-s(x-\theta)^2}$) around an optimal expression of $\theta = 0.5$, for two strengths of selection ($s = 10$, the default simulations, and $s = 50$, the strong selection setting).

#### Supplementary Figure S2.

| | Before domestication | After domestication | | |
|---|---|---|---|---|
| | | Plastic | Stable | Non-selected |
| Plastic (p) | 6 | 2 | 2 | 2 |
| Stable (s) | 6 | 0 | 4 | 2 |
| Non-selected (n) | 12 | 0 | 4 | 8 |
| Total | 24 | 2 | 10 | 12 |

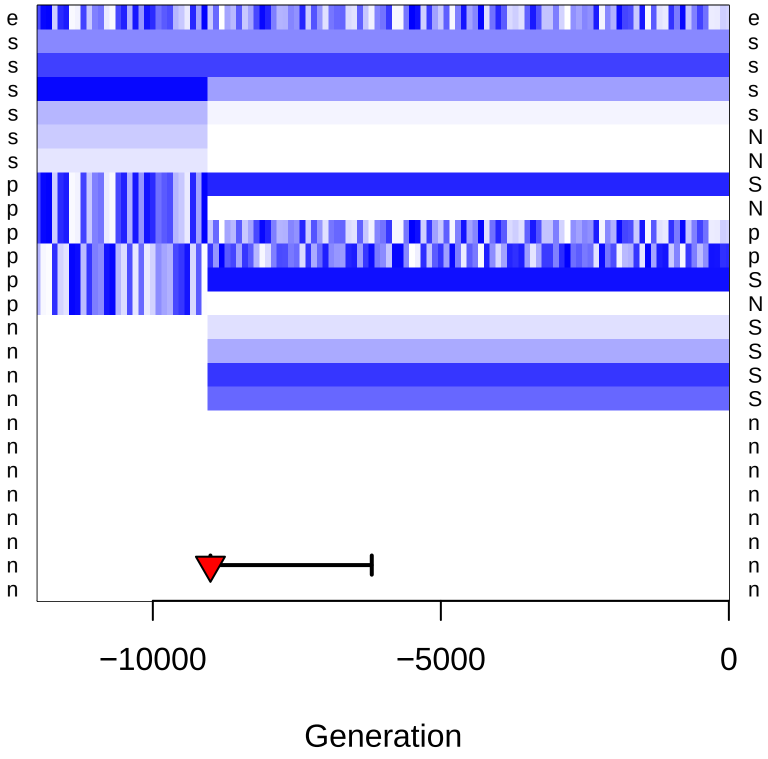


**Description of the selection switch associated with domestication.** Top: summary table of the change in selection regime for the 24 genes of the network. Three categories of genes were considered. Stable (s) genes have a constant optimum. For half of them, the optimum changed once, at the onset of domestication (the other half keep the same optimum throughout the whole simulation). For Plastic (p) genes, the optimum tracks the environmental signal every generation (with a +1 correlation for half the genes, and a -1 correlation for the other half). The expression of non-selected genes (n) does not affect the fitness function (in practice, the corresponding selection coefficient is set to 0). The total number of selected genes (12 out of 24) remains the same before and after domestication. Bottom: illustration of the variation of gene expression optima from a single simulation. The figure indicates the expression of the sensor gene (first line, e), and the optimal expression for all other genes (light: low expression, dark: strong expression). White stands for unselected genes, and capital letters for genes which status changed.

#### Supplementary Figure S3.

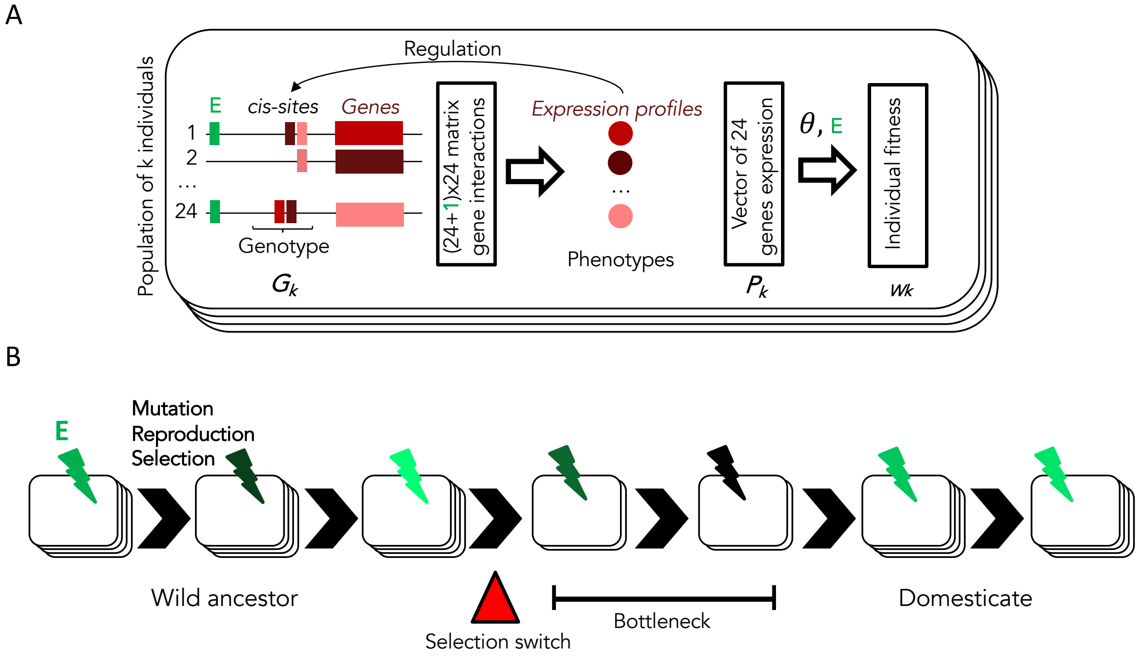


**Evolution of simulated networks.** A: At the individual ($k$) level, we simulated a network of 24 genes (transcription factors) regulated in cis (cis-sites). Allelic states at these sites constitute the genotype of an individual ($G_k$). Together with an environmental cue ($E$) that regulate some of the genes (plastic genes), those sites determine a matrix of gene interactions that determines the phenotype of $k$ ($P_k$), a vector of expression profiles for each of the 24 genes. Individual fitness ($w_k$) in a multivariate landscape is determined by the distance to the selection target ($\theta$) for 12 selected genes, $\theta$ being constant for stable genes, and variable (correlated to the environment $E$) for "plastic" genes. Unstable gene networks (unable to achieve a stable equilibrium in gene expression) are associated with a penalty in the fitness function. B: At the population level, $k$ individuals are submitted to mutations of cis-sites that affect $G_k$, as well as reproduction and selection at each generation in a fluctuating environment ($E$). At the time of domestication, the selection switch modifies the selection target and mimicks evolution towards phenotypic stability while the bottleneck translates into a shrink in population size.

### Supplementary Figure S4.

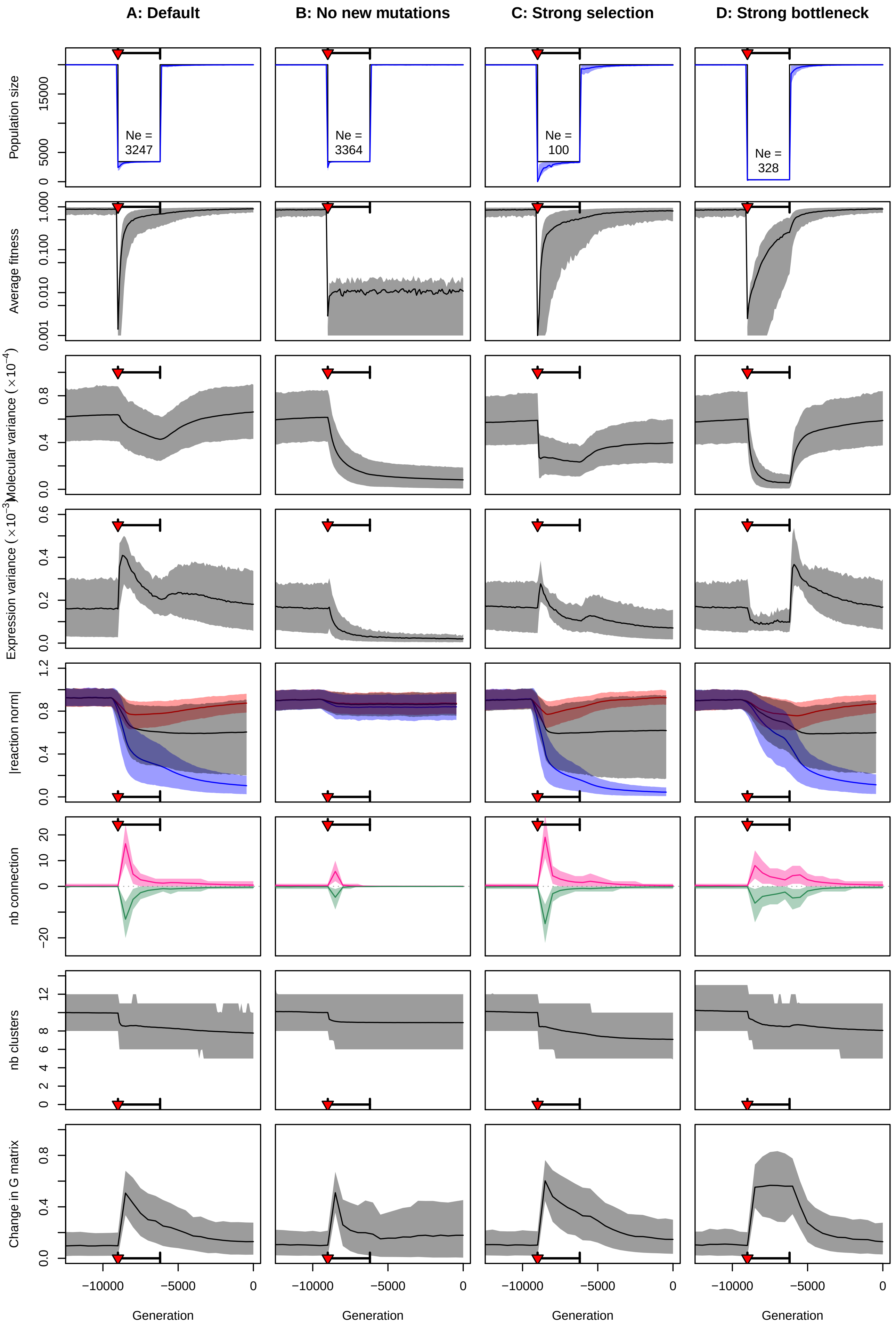

**Sensitivity of the simulations to mutational and demographic parameters.** The left column (A) stands for the default parameter set (maize domestication scenario). B: No new mutations: the response to domestication relied only on the standing genetic variation (mutation rate set to 0 at the onset of domestication); C: Strong selection: the selection coefficient was set to 50 instead of 10 (as illustrated in Figure S1); D: Strong bottleneck: the strength of the bottleneck was 10 times stronger (350 individuals instead of 3500). Rows stand, respectively, for:

1. Census and effective population sizes. Census sizes are model parameters, effective population sizes $N_e$ are estimated from the variance in fitness (as detailed in the methods section). Effective population sizes indicated in the figure were computed as the harmonic mean $N_e$ over the whole bottleneck.

2. Mean fitness. Thick line: Average fitness across 1000 replicates; grayed area: 10% and 90% quantiles of the mean fitness over 1000 replicates.

3. Molecular variance (averaged over the whole network), 10% and 90% quantiles. See Fig 4A for details.

4. Expression variance (averaged over the whole network), 10% and 90% quantiles). See Fig 4B for details.

5. Reaction norms of genes that were selected to be plastic before domestication (as in Fig. 3B). Red: plastic genes after domestication, Black: neutral genes after domestication, Blue: stable genes after domestication. Thick lines stand to the average, and the colored area for the 10% - 90% quantiles.

6. Number of gained and lost connections (same color code as in Fig. 6A).

7. Number of clusters (as in Fig. 6B), thick lines stand for the mean and the colored area for the 10% - 90% quantiles.

8. Evolution rate of the G matrix (distance between successive genetic correlation matrices, as described in the methods section and in Fig. S6A). Thick lines stand for means, grayed areas for 10% - 90% quantiles.

#### Supplementary Figure S5.

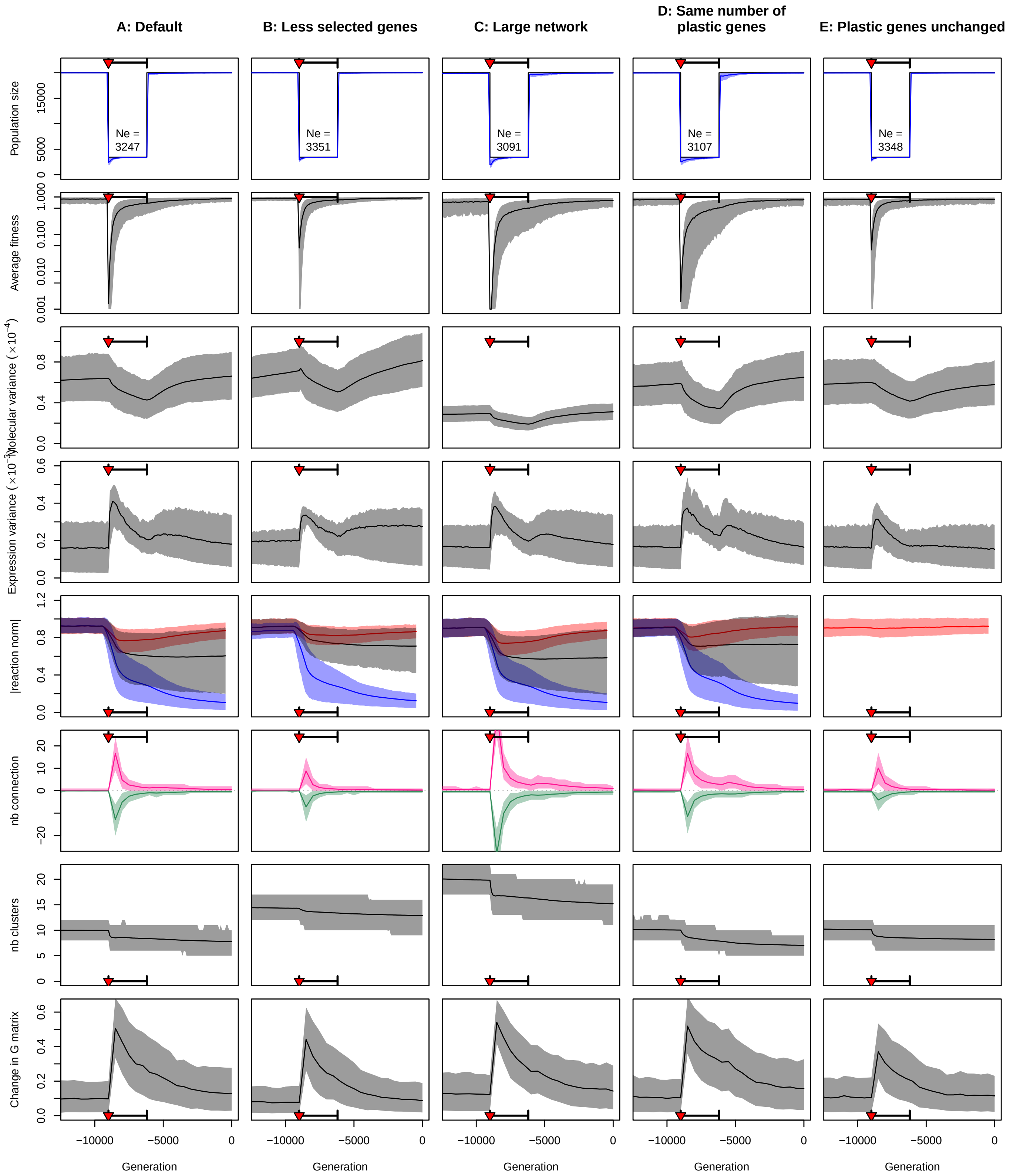


**Sensitivity to variations in selection and gene network parameters**. The left column (A) represents the default parameter set (maize scenario). B: Less selected genes: the network still contained 24 genes, but only 6 (instead of 12) evolved under selection, with the same proportion of plastic, stable, and non-selected genes; C: Large network: 48 genes (+ environment) were considered in the simulation, with 24 genes under selection. D: Same number of plastic genes: Domestication was not associated with less selection on plasticity; 6 genes are still selected for plasticity during the domestication process, but these were not the same as the plastic genes before domestication. E: Plastic genes unchanged: the six plastic genes after domestication were the same as the ones before domestication. Rows are the same as in Fig S4.

### Supplementary Figure S6.

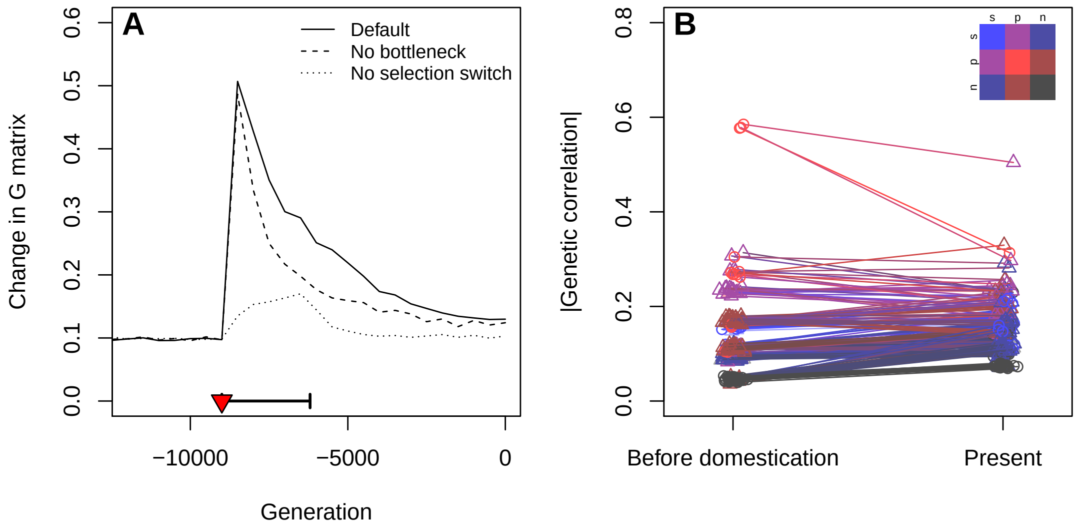


**Evolution of the genetic coexpression structure.** (A) The evolution of the genetic covariance matrix G of all gene expressions was tracked by measuring the distance between G every 500 generations (details in the methods section). (B) Genetic correlations before (generation -9000) and after (generation 0) domestication were compared between all pairs of genes. The color code indicates the category of the genes compared: blue for stable/constant, red for plastic, black for non-selected, and intermediate colors for comparisons between genes of different categories. Circles denotes correlations between genes from the same category, triangles from different categories. The x-axis was slightly jittered for clarity.

#### Supplementary Figure S7.

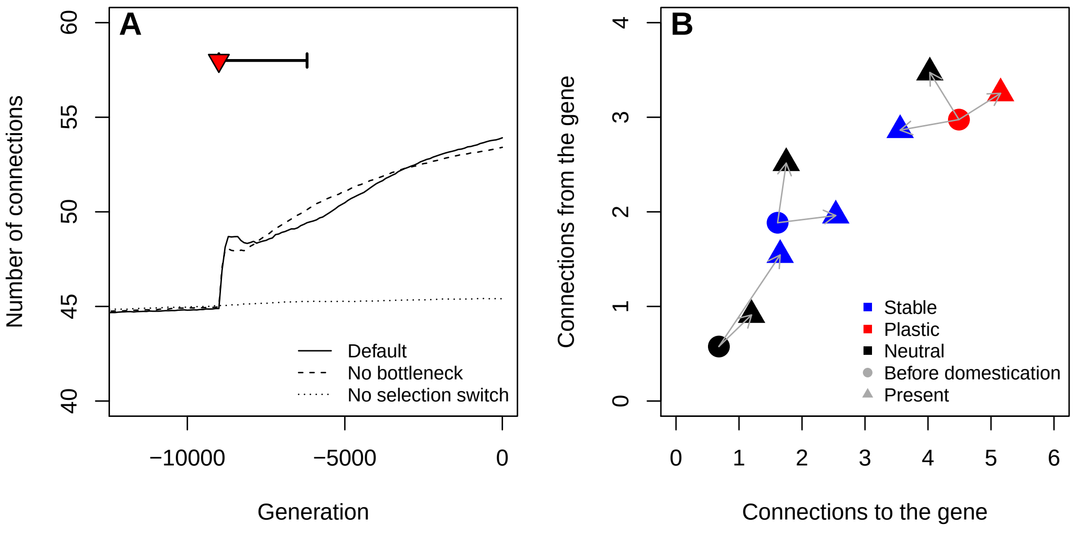


**Evolution of the connections in the newtork.** A: Evolution of the average number of strong connections in different evolutionary scenarios. B: Evolution of the numbers of connections to (number of regulators of the target gene) and from (number of genes regulated by the target gene) each type of gene during domestication in the default simulation (bottleneck and selection switch).

### Supplementary Figure S8.

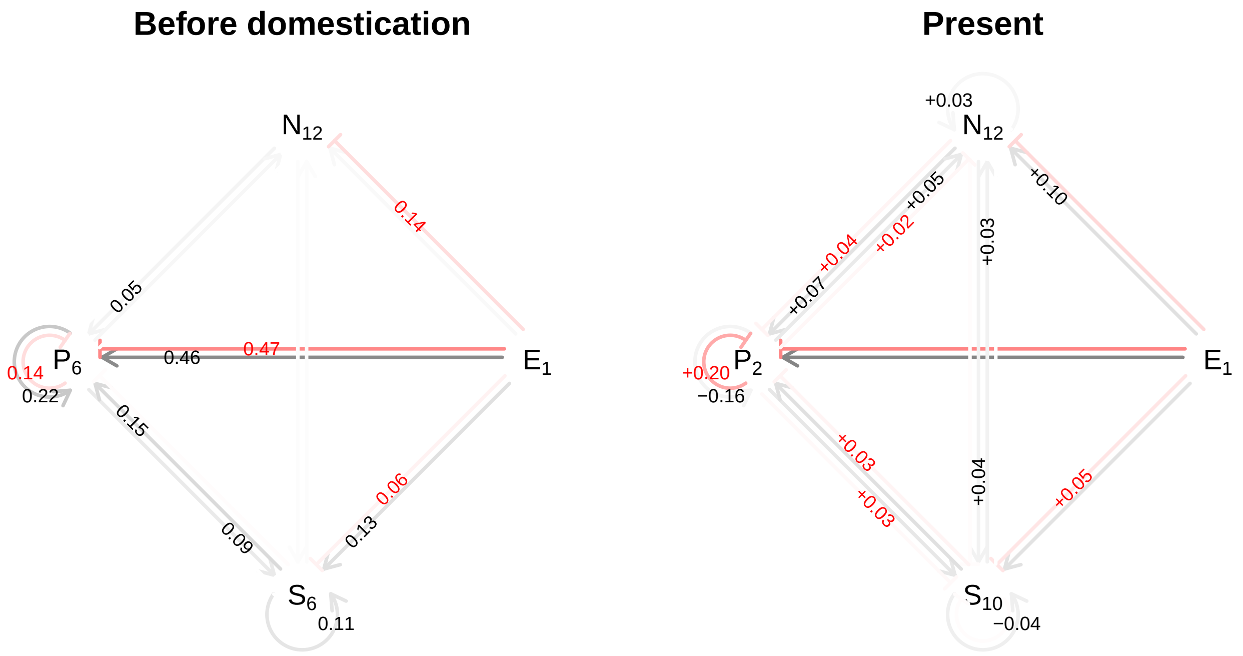


**Summary of the topological changes in the network.** The figure displays the location of the strong connections in the network, based on the effect of regulation on gene expression (details in the methods section). Four categories of genes are represented: non-selected (N), plastic (P), stable (S); (E) stands for the gene that is directly reflecting the environment. Subscripts indicate the number of genes for each category. The left panel indicates the frequency of connections between each gene category (i.e. the number of strong connections divided by the theoretical maximum) before domestication. Right: Network at the end of the simulations. Numbers stand for the differences compared to the left panel. In both panels, connections between genes from the same category are indicated as circular arrows. Red arrows stand for negative (inhibition) effects, black arrows for positive (activation) effects; the color intensity is proportional to the connection frequencies. Connection frequencies below 5% were omitted for clarity.

### Supplementary Figure S9.

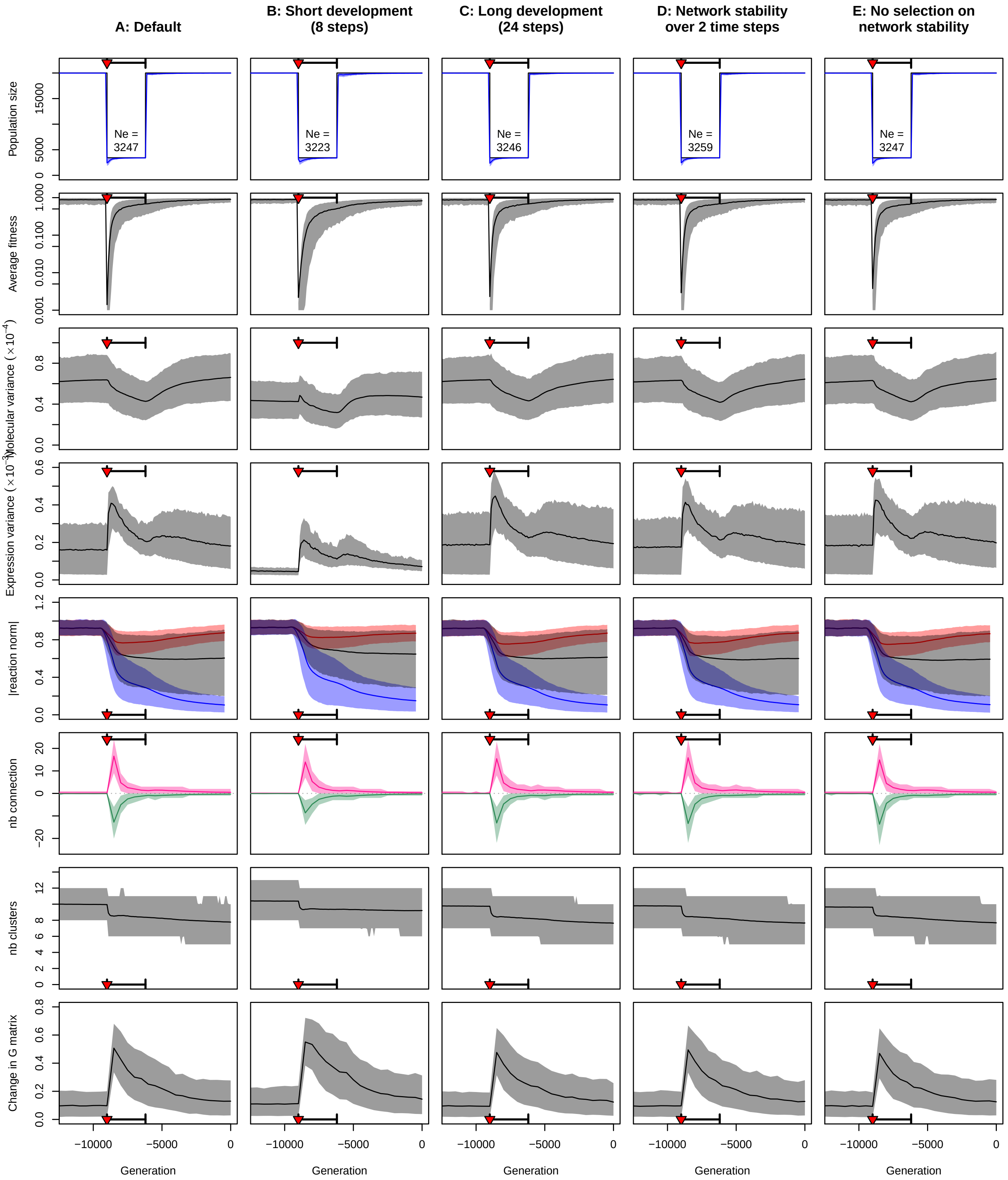


**Sensitivity of simulation results to arbitrary choices in the gene network model dynamics.** A: Default simulations (maize scenario). B: Short development (8 time steps instead of 16 for the Default case). C: Long development (24 time steps instead of 16). D: The network stability was assessed over the two last time steps (instead of 4). E: there was no selection on network stability. Rows stand for the same indicators as in Fig S4.
